# Bioremediation of hydrocarbon contaminated soil from Carlini Station, Antarctica: effectiveness of different nutrient sources as biostimulation agents

**DOI:** 10.1101/753384

**Authors:** Julia Villalba Primitz, Susana Vázquez, Lucas Ruberto, Alfredo Lo Balbo, Walter Mac Cormack

## Abstract

Logistics and scientific activities carried out in Antarctic stations entail the risk of contamination by fuels. Among remediation strategies, biostimulation of chronically contaminated Antarctic soils significantly improves the efficiency of hydrocarbons (HCs) removal. The aim of this study was to evaluate the performance of different nutrient formulations as biostimulation agents, in order to improve the elimination of diesel fuel from Antarctic soils, in both oxic and anoxic conditions. A field test was performed in microcosms (15 kg of soil each) as experimental systems. Each microcosm was prepared by triplicate, sampled every 10 days over a 50-days period and sampled again one year later. Changes in bacterial communities, and qualitative and quantitative HCs analysis were determined. Our results showed that, during the early stages of the process, a multi-component commercial product like OSEII^®^ (containing nutrients, enzymes and surfactants) determines a rapid elimination of HCs with changes in the structure of the bacterial soil community, whereas a more cost-effective slow-release fertilizer like Nitrofoska^®^ would be efficient in a long-term bioremediation process.

## 1. Introduction

Due to its low cost, effectiveness, versatility, and safety of use and storage compared to other available options, fuels derived from crude oil are, still today, the most used energy source. These fuels represent an important fraction of the international market (Zhong et al., 2016) and their use at global scale increases the risk of spills. The presence of hydrocarbons in natural environments represents an environmental problem throughout the world, including polar and isolated regions such as the Arctic (Kachinskii et al., 2014, Bennett et al., 2015) and Antarctica (Szopińska et al., 2016, Raymond et al., 2017, Vázquez et al., 2017), where hydrocarbons pollute soils, sediments, and water, mainly around human settlements (Hughes and Stallwood, 2006; Braun et al., 2014; de Jesus et al., 2015).

Most of the nearly 50 Antarctic stations using fuels derived from crude oil to generate energy and heat are settled on the few ice-free regions (less than 0.3% of the continent’s total area), where most of the terrestrial biota is also established (Aislabie et al., 2004). To protect these sensitive ecosystems, regulations related to the preservation of the environment were established in the Antarctic Treaty Protocol, signed in 1991 (https://www.ats.aq/e/ats.htm). For this reason, the development of site-specific methods for cleaning Antarctic soils polluted with hydrocarbons is required. These methods should be environmentally friendly, simple, inexpensive, and effective at low temperatures. In addition, to meet environmental policies suggested by the Antarctic Treaty and its Protocol on Environmental Protection, they must involve exclusively the use of autochthonous organisms.

Natural attenuation comprises a variety of physical, chemical, and biological processes and involves autochthonous microorganisms acting without human intervention (EPA, 1999). Although this strategy is considered an effective tool for bioremediation of some soils and groundwaters (Mulligan and Yong, 2004); low temperatures, low evaporation rates, viscosity of long-chain hydrocarbons, and lower water solubility of the contaminants convert natural attenuation in a poorly efficient process for removal of hydrocarbons from soils in cold areas (Camenzuli and Freidman, 2015). In Antarctica, most of the human settlements are concentrated in the coastal areas of the West Antarctic Peninsula (WAP), where soil temperatures reach values above the freezing point only for about three months, while the soils stay frozen and covered by ice and snow for the remaining nine months of the year. These conditions also limit natural attenuation, making it an ineffective strategy to reach acceptable levels of contaminants in soils (Dias et al., 2012). For this reason, hydrocarbon removal from WAP soils mostly requires designing and application of active bioremediation strategies.

*In situ* (or, alternatively, *on-site*) bioremediation, involving the activity of a wide array of microorganisms is considered the best approach to reach the maximum removal of hydrocarbons from soil (Goswami et al., 2018). Our previous studies with Antarctic soils showed that biostimulation with N and P significantly improves removal of aliphatic hydrocarbons from contaminated soils (Ruberto et al., 2009; Dias et al., 2015; Martínez Álvarez et al., 2017), whereas bioaugmentation does not seem to provide additional advantages when the soil has a long-term exposure to fuels and the indigenous microbiota is well adapted to their presence in the environment (Vázquez et al., 2009; Ruberto et al., 2010). Similar results obtained with different soils and under different climates were reported elsewhere (Kauppi et al., 2011; Zawierucha and Malina, 2011; Abed et al., 2014; Wu et al., 2016).

A successful biostimulation depends on both the type and the amount of N and P sources, and several studies have focused on the optimization of the amount of nutrients to add to the soil to balance the high C input from the contaminant hydrocarbons. In this regard, low hydrocarbon biodegradation rates have been related to insufficient amounts of nutrients added as fertilizers (Huesemann, 2004), whereas an excess of nutrients could inhibit the degradation of hydrocarbons (Ruberto et al., 2003; Grace Liu et al., 2011). Therefore, to keep a right balance of nutrients throughout a bioremediation process is one of the main factors to deal with for a successful treatment. However, the optimal C:N:P ratio reported for hydrocarbon degradation in soils is widely variable (Lee et al., 2007), making it necessary to carry out optimization tests for each particular soil and situation. A study by Emami et al. (2014) evidenced that the nature of the fertilizer also has an important role in oil degradation. In this sense, many different N and P sources have been tested for bioremediation. Inorganic salts (i.e. ammonia, nitrate, and phosphate) have been reported as simple and efficient nutrient sources to sustain hydrocarbon biodegradation (Grace Liu et al., 2011; Martínez Álvarez et al., 2017). However, due to their high water solubility, they are lost by runoff, being difficult to control their dissolution rate and maintain an adequate concentration to support microbial growth with a minimal risk of toxicity in the early stages after the addition. Therefore, for the remediation of hydrocarbons alternative sources of nutrients with a better cost/benefit ratio have been tested. Among these sources, we can mention urea (Simpanen et al., 2016); various biowastes like soy cake (Dadrasnia and Agamuthu, 2014), poultry droppings (Ezenne et al., 2014), oil palm empty fruit bunch, sugarcane bagasse (Hamzah et al., 2014), fish meal (Dias et al., 2015); NPK inorganic fertilizers (Silva-Castro et al., 2015) and other commercial bioremediation agents (Dias et al., 2012). For each particular case, the selection of the best source of nutrients seems to depend on the characteristics of the soil under treatment, because the microbiota thriving in each soil is different and can exhibit dissimilar metabolic requirements. Therefore, as it was stated by Rittmann et al. (2006), understanding the ecology that drives the structure of bacterial communities is crucial for the development of any environmental biotechnology process. In this sense, getting knowledge about the structure and dynamics of microbial communities during a bioremediation process offers perspectives to evaluate the performance of the applied strategy.

Concerning oxygen availability, the degradation of hydrocarbons in soils under aerobic conditions proved to be effective, and frequently leads to a high-rate removal. However, in certain remote locations like the Antarctic stations, aeration of huge amounts of soil is expensive and requires a complicated logistics. In addition, precipitation and ice covers cause the soil to become anaerobic, and even if it is aerated, transient anaerobic pockets form in the soil matrix due to physical, chemical or biological factors. Under these situations, microbes able to utilize alternative electron acceptors like nitrate, sulfate or fumarate could be an interesting alternative for the anaerobic degradation of hydrocarbons at contaminated sites (Sampaio et al., 2017).

Considering all the above stated, the aim of this study was to evaluate the performance of different biostimulation agents on the removal of diesel hydrocarbons from Antarctic soils under both oxic and anoxic conditions, using microcosms as experimental systems. Heterotrophic and hydrocarbo*n*-degrading viable bacterial counts and DGGE fingerprints were used to monitor variations in bacterial numbers and infer qualitative changes in bacterial populations throughout a field-scale bioremediation experiment.

## 2. Materials and Methods

### 2.1. Study area and experimental design

The experiment was carried out at Carlini Station (62° 14’S, 58° 40’ W), located on Potter Peninsula, Isla 25 de Mayo (King George Island), South Shetland Islands, Antarctica (Fig.1A). The soil used in this study was collected near the main fuel storage tanks and was chronically contaminated with diesel (Antarctic gasoil: AGO), but also exposed to fresh contamination caused by frequent leaks from the connection pipes and the refueling activities. After collection, large stones (> 1cm diameter) were removed manually and the soil was sieved (10 mm mesh), vigorously mixed to homogeneity and finally disposed in plastic containers simulating small biopiles with 15 kg of soil each (Fig.1B).

**Fig. 1.**
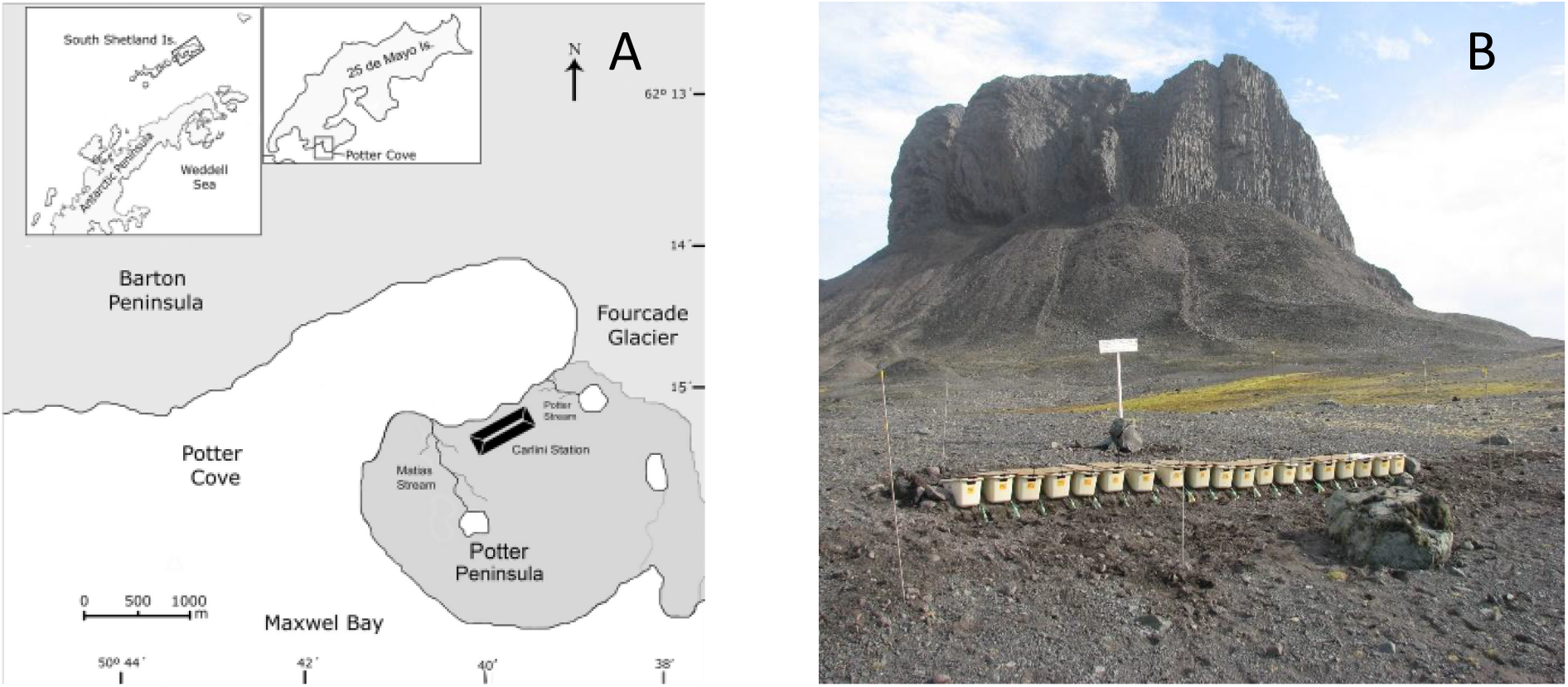
A) Geographic location of Potter Cove and Potter Peninsula showing Carlini Station, where the microcosms were located. B) Microcosms were kept outdoors in an area with no traffic and exposed to environmental conditions.

Treatments with different nutrient sources were performed in triplicate and the microcosms were distributed on the ground, covered by plastic lids as protection against wind, rain, and snow (Fig. 1B, S1). Based on the total organic C estimated from the hydrocarbon content (7620 ± 680 mg AGO/kg_dw_ soil), IN and ANA microcosms were biostimulated with inorganic salts (NH_4_NO_3_ 0.17% w/w, Na_2_HPO_4_ 0.027% w/w) to achieve a C:N:P ratio of 100:10:1. Additional amounts of nutrients (corresponding to 20% of that added at the time of preparation of the microcosms, day-0) were added after each sampling. To evaluate the effect of anoxic conditions on the degradation of hydrocarbons, IN was weekly aerated by mixing, whereas ANA was not mixed at all. NPK microcosms were designed to favor a more constant availability of nutrients. These microcosms were aerated by mixing weekly and supplied with a commercial slow-release granular fertilizer, Nitrofoska^®^ (0.5% w/w), which was added in fractions: 50% at day-0 and 50% after 10 days. CP microcosms were supplemented with a multi-enzymatic commercial product, OSEII^®^, Oil Spill Eater International Corp., a bioremediation agent listed on the US Environmental Protection Agency’s National Contingency Plan for Oil Spills and reported as a biological enzyme additive which contains nutrients, surfactants, and vitamins to promote microbial growth. According to the manufacturer’s instructions, 15 ml of OSEII^®^ were added per kg of wet soil at day 0 and the microcosms were weekly aerated by mixing. Two control treatments were also performed: i) AC, abiotic control, a microcosm to evaluate abiotic (non-biological) loss of hydrocarbons, in which the soil was poisoned with 0.3% w/w HgCl_2_, and ii) CC, community control, non-fertilized microcosms with weekly aeration by mixing, to evaluate the removal of hydrocarbons by the indigenous microbiota.

All experimental units were kept outdoors for one year, under natural environmental conditions, although covered with a plastic lid to protect from flooding and snow filling and prevent excessive washing of nutrients and hydrocarbons. Sampling was performed during the austral summer, at days 0, 10, 20, 30, 40, 50 and 365. Samples were composed of three portions taken randomly from each microcosm (Fig. S2) and were frozen at −20°C until processed, except for bacterial counts and pH measurements which were carried out immediately in the laboratory at Carlini Station.

### 2.2. Bacterial counts

Viable cell counts were measured in each sample to estimate the number of cultivable heterotrophic aerobic bacteria (HAB) and hydrocarbo*n*-degrading bacteria (HDB). One gram of soil was suspended in 10 ml of saline solution containing Tween20 (0,1%) and te*n*-fold serial dilutions in saline solution were prepared from the suspension. Then, 100 μl of each dilution were plated in R2A agar (for HAB counts) or in solid saline basal medium with AGO as the sole carbon and energy source (for HDB counts). Plates were incubated at 10 °C for one week (HAB) or three weeks (HDB). Results were expressed as colony-forming units per gram (dry weight) of soil (CFU/g_dw_) and the significance of the differences between mean values was calculated using one-way ANOVA and *post-hoc* Tukey’s test.

### 2.3. Analysis of hydrocarbons

The qualitative patterns of hydrocarbons by gas chromatography (GC-FID) using dichloromethane (HPLC grade, Sintorgan) as solvent, and the quantification by infrared spectrophotometry (IR) were performed as described in Martínez Álvarez (2017). Three subsamples from each composite sample were extracted and measured independently. Total petroleum hydrocarbons (TPH) content was expressed in mg AGO/kg_dw_ and calculated as the average of the triplicate subsamples. Hydrocarbons removal efficiency at day-50 and day-365 was calculated from the difference of TPH at those sampling times and the value at the initial time. The significance of the differences between treatments was calculated using one-way ANOVA and *post-hoc* Dunnett test.

### 2.4. Extraction of total DNA and PCR amplification for DGGE

For bacterial communities’ fingerprinting, total DNA was extracted from 0.5-1 g of well-homogenized soil using the Power Lyzer Power Soil DNA Isolation Kit™ (Mobio), following the manufacturer’s instructions with some modifications: after addition of Bead Solution, 20 μl lysozyme (300 mg/ml) were added and the tubes were incubated for 30 min at 37°C. After that, samples were subjected to three cycles of freezing (−80°C) and thawing (65°C). Three independent PCR amplifications were done from each sample. Fragments of the V3 hypervariable region of the bacterial 16S rRNA gene were amplified using the GC-341F and 534R primers (Muyzer et al., 1993). PCR reactions contained 40 ng of total DNA, 0.5 μM of each primer, 0.2 mM of each deoxyribonucleotide triphosphate, 0.25 mg/ml of bovine serum albumin, 1.5 U GoTaq™ (Promega) and 1X GoTaq™ buffer, in a final volume of 50 μl. PCR amplification consisted of an initial denaturation step for 5 min at 94°C; 35 cycles consisting of 94 °C for 45 sec; 55 °C for 45 sec; 72 °C for 45 sec, followed by a final extension at 72 °C for 5 min. The quality and quantity of extracted DNA and PCR products were analyzed by agarose gels electrophoresis (1.5% w/v) stained with GelRed™ (Biotium, Inc.).

### 2.5. Denaturing gradient gel electrophoresis (DGGE)

Differences in the structure of the bacterial communities present in all samples were inferred by DGGE fingerprinting. Total DNA was extracted from each sample and used as template in 3 independent PCR reactions, which were then pooled and concentrated to 35 μl using a Vacuum Concentrator (Hetovac VR-1). About 500 μg DNA from each sample were loaded into gels containing 8% (w/v) polyacrylamide (acrylamide:N,N’-methylene bisacrylamide, 37.5:1) in 1x TAE buffer (10 mM sodium acetate, 0.5 mM Na_2_-EDTA and 20 mM Tris, pH 7.4), with a denaturing gradient ranging from 45% to 60% (with 100% denaturant containing 7M Urea and 40% v/v formamide). DGGE was performed in a TV400-DGGE system (Scie-Plas Ltd.). Gels were run at 65°C for 16 h at 60V, stained for 1 h with SyberGold™ (Invitrogen) in 1X TAE buffer and documented under UV light. DGGE profiles were compared based on the presence or absence of bands using the Jaccard similarity coefficient and dendrograms were constructed according to the unweighted pair group method with average means (UPGMA) using the Gel Compar II™ software package (Applied Maths).

## 3. Results

### 3.1. Bacterial counts

Biological activity in the different microcosms was inferred from the changes observed in the cultivable fractions of HAB and HDB (Fig. 2). No colonies grew from the AC microcosms, confirming an efficient poisoning of the soil microbiota by the mercuric chloride. As a general trend, HAB counts increased in all microcosms throughout the study (Fig.2A). The high counts obtained at day-0 from the non-fertilized CC microcosms proved the positive effect of mixing and aeration on bacterial growth. However, the counts decreased after 30 days, to end at day-50 and day-365 with lower HAB counts than those of the biostimulated treatments. CP microcosms had an early increase in HAB counts that remained higher than those recorded for any other treatment until the end of the experiment. On the other hand, HAB evolved differently in NPK microcosms, with no significant change until day-20 followed by a slight but continuous increase until day-50. After one year, at the end of the experiment, the HAB counts stayed like on day-50 and were almost equal for all treatments, except for CC, where HAB counts were significantly lower (*p*<*0.05*; except NPK) (Fig.2A).

**Fig. 2.**
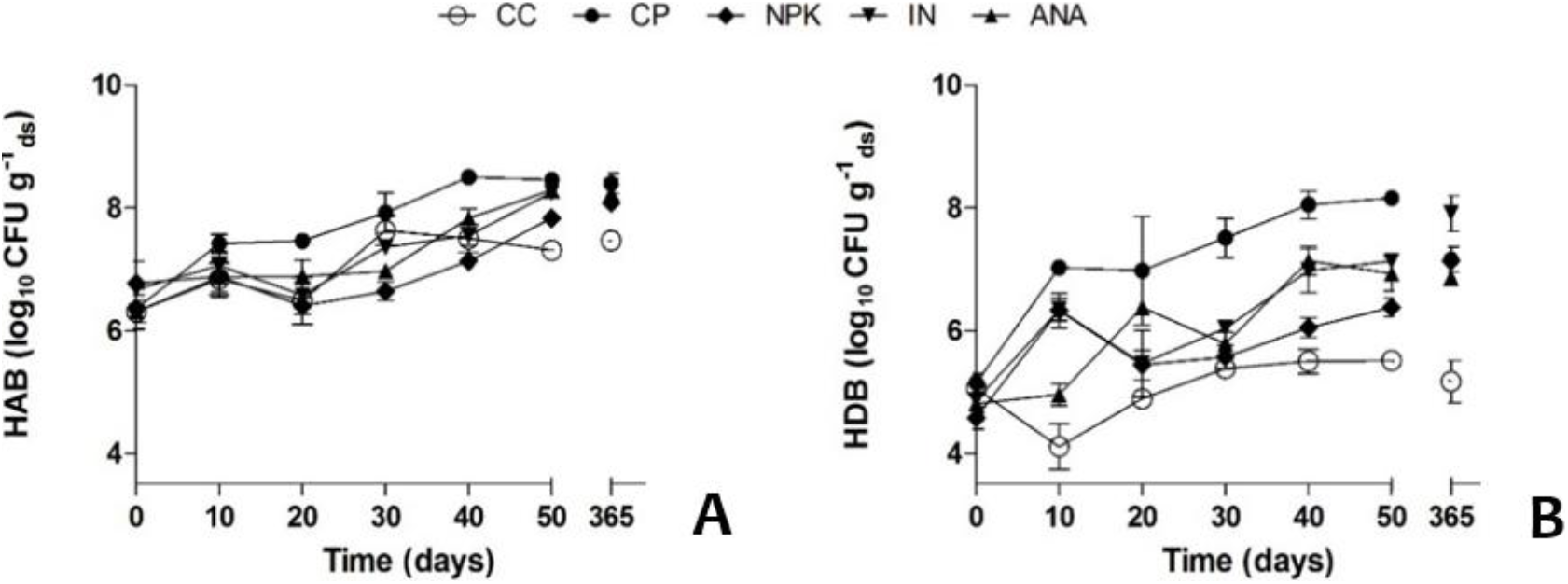
Total cultivable heterotrophic aerobic (A) and hydrocarbon degrading (B) bacterial counts recorded for all microcosms. CC: non-fertilized control, IN: biostimulation with inorganics salts, ANA: biostimulation with inorganic salts without mixing, NPK: biostimulation with slow-release granular fertilizer, CP: biostimulation with a commercial bioremediation agent. Symbols represent the mean of three independent replicate microcosms for each treatment. Error bars indicate SD.

HDB counts at the initial time ranged between 3.97×10^4^ and 1.46×10^5^ CFU/g_dw_ (Fig. 2B). In CC microcosms, HDB counts dropped by day-10 and then increased, but remained below the HDB counts of the other treatments at any sampling time, and reached significantly lower counts towards day-50 (*p*<*0.05*). Biostimulation favored the cultivable fraction of HDB, evidenced by the significantly higher (*p*<*0.05*) counts recorded for all treatments compared to the CC control, confirming that the addition of nutrients is indispensable to promote the growth of the HBD in this soil. Moreover, the HDB counts at day-50 in CP, IN, ANA and NPK treatments were significantly higher (*p*<*0.05*) than those in CC. As was observed with HAB counts, HDB in CP soon increased to reach values close to10^8^ UFC/g_dw_ at day-50, approximately one order of magnitude higher than in the other fertilized microcosms. HDB in NPK microcosms showed a similar trend to HAB counts, with values always lower than those observed for the other treatments, likely due to the slow release of N and P mentioned above. Finally, after one year, all the fertilized microcosms reached HDB counts between 7.4×10^6^ and 9.14×10^7^ CFU/g_dw_, all values significantly higher (*p*<*0.05*) than those recorded from CC microcosm (1.18×10^5^ CFU/g_dw_).

### 3.2. Efficiency of the biostimulation agents for removal of hydrocarbons from soil

The hydrocarbon removal efficiency of each treatment at day-50 and day-365 is shown in Figure 3. After 50 days, no significant differences were detected between the controls, AC (12 ± 3%) and CC (18 ± 3%). Concerning the biostimulated treatments, all but ANA showed significantly higher hydrocarbon removal efficiencies (*p*<*0.05*) at day-50, compared to AC. The microcosms added with the slow-release granular fertilizer (NPK) and the commercial agent (CP) were considerably more efficient (32 ± 6% and 50 ± 9%, respectively) than the non-fertilized control CC (*p*<*0.05*) after 50 days of treatment (Fig.3A). The higher removal efficiency in the microcosms added with the commercial product (CP) was in accordance with the early increase observed in bacterial counts.

**Fig. 3.**
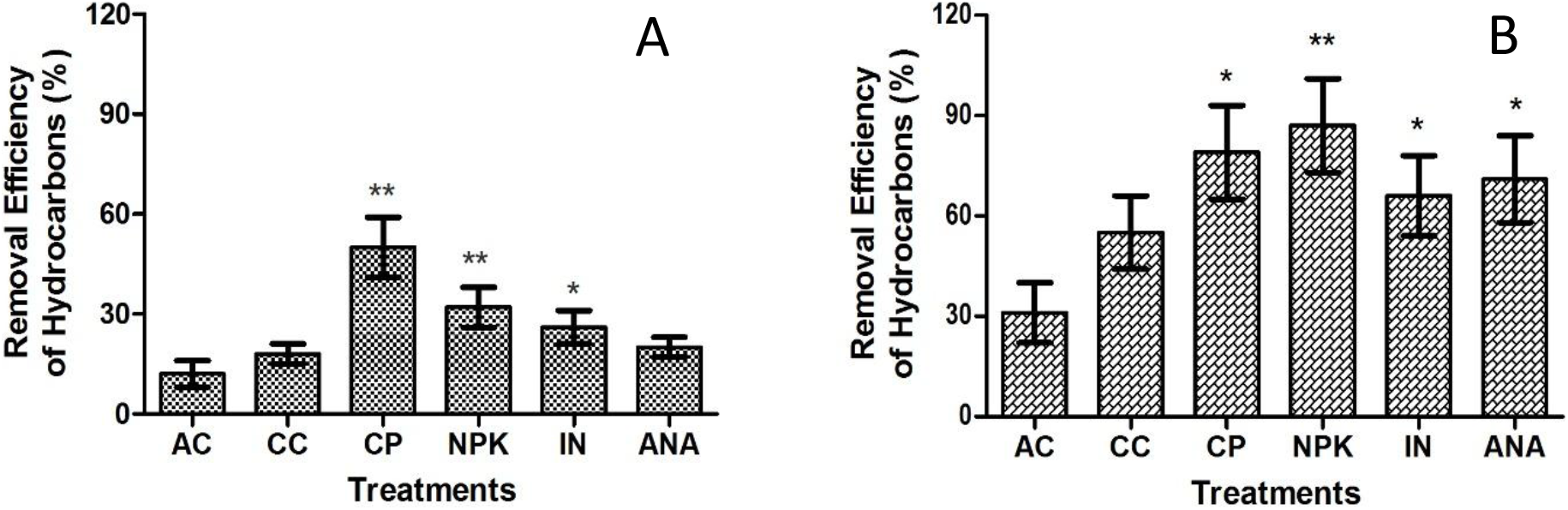
Hydrocarbons removal efficiency (%) at T5, day-50 (A) and T6, day-365 (B) of the experimental treatments applied using different biostimulation strategies during the bioremediation of an Antarctic soil chronically exposed to diesel fuel. AC: abiotic control, CC: non-fertilized control, IN: biostimulation with inorganics salts, ANA: biostimulation with inorganic salts without mixing, NPK: biostimulation with slow-release granular fertilizer, CP: biostimulation with a commercial bioremediation agent. (*) indicates significant differences compared to AC and (**) indicates significant differences compared to AC and CC (*p*<*0.05*).

After one year *on site*, all microcosms with biological activity had lower hydrocarbon contents than the abiotic control, which suggests that biodegradation contributed to the total elimination of hydrocarbons from the soil, in addition to abiotic loss (Fig.3B). CP, NPK, IN and ANA treatments achieved hydrocarbon removal efficiencies of 79%; 87%; 66% and 71%, respectively, all significantly higher values (*p*<*0.05*) than the 31% recorded for the abiotic control AC. Only NPK treatment reached values significantly higher (*p*<*0.05*) than the 55% recorded for the non-fertilized control CC.

Table 1 shows the ratios between the concentrations of some individual hydrocarbons. In AC, *n*-C10-13/*i*-C16 ratios decreased during the first 50 days of the experiment, with *n*-C14-15/*i*-C16 and *i*-C13/*i*-C16 almost unaltered, suggesting a small loss by evaporation of the lighter compounds (up to *n*-C13), which was greater after one year and more intense the shorter the length of the *n*-alkane chain (Fig. 4 A-B-C). Considering the same ratios, no major changes were observed on day-0 between CC and the most efficient treatments (NPK and CP) compared to AC, except for *n*-C10/*i*-C16. This suggests that *n*-C10 could have been degraded to some extent by the microbiota stimulated by the aeration produced in the soil by mixing, shortly after the preparation of the microcosms. In CP, the same was observed for *n*-C10 and, to a lesser extent, also for *n*-C11-13, probably due to the stimulation of the microbiota by the rapid action of OSEII^®^ that improved the bioavailability of hydrocarbons and the supply of nutrients. At T5, all ratios in CC and AC were rather similar with no substantial changes in the chromatographic pattern of residual hydrocarbons in CC, compared to the initial time (Fig. 4 D-E). NPK had *n*-C10-13/*i*-C16 values lower than AC, which were even lower in CP, as was the *i*-C13/*i*-C16 ratio, involving less biodegradable compounds with a different rate of evaporation (Fig. 4 G-H, J-K). After one year, these ratios in NPK and CP, were much lower even than in the non-biostimulated control CC, indicating that the hydrocarbons were not only removed by washing and evaporation but also by biodegradation, especially the lighter compounds (up to *i*-C13). Contrary to the fuel used in the Australian Antarctic Stations (SAB), in the diesel used by Argentina in Antarctica (AGO), *i*-C13 is more abundant than *i*-C14 (Fig. S3). Therefore, and since *i*-C13 has shorter chain length and is more volatile, when the ratio *i*-C13/*i*-C14 becomes less than 1 there is indication of evaporation alone or accompanied by advanced degradation. In AC and CC controls, this slightly occurred only after one year (Fig. 4 B-C, E-F), but was more pronounced in NPK (Fig. 4 H-I) and even faster (already at day-50) in CP microcosms (Fig. 4 K-L), confirming the biological degradation promoted by biostimulation in those microcosms.

**Table 1:**
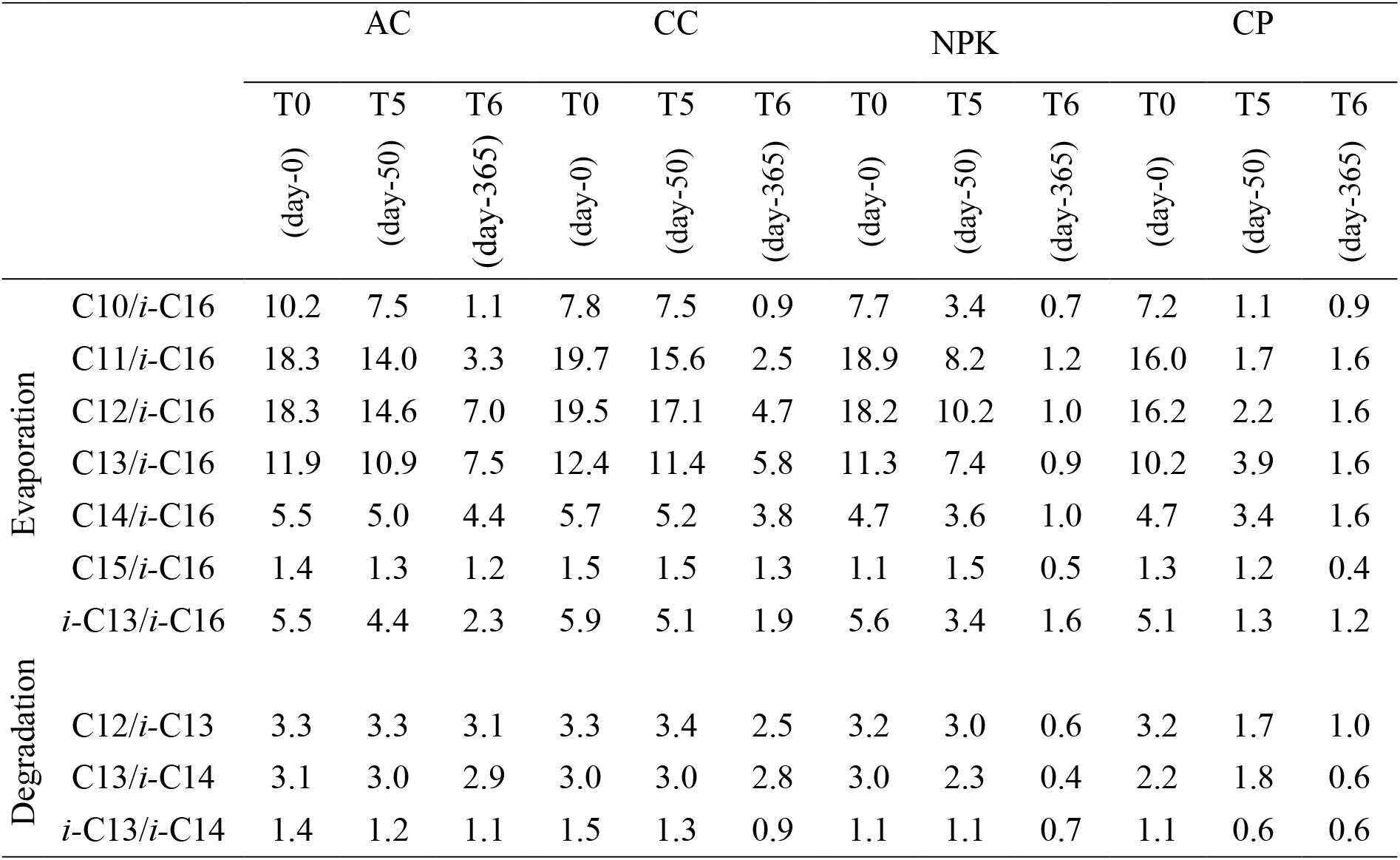
Ratios between the concentrations of some individual hydrocarbons in soil samples from the most efficient treatments (CP and NPK), the non-fertilized control (CC) and the abiotic control (AC) at the beginning of the experiment (T0) and after 50 (T5) and 365 (T6) days of treatment.

**Fig. 4.**
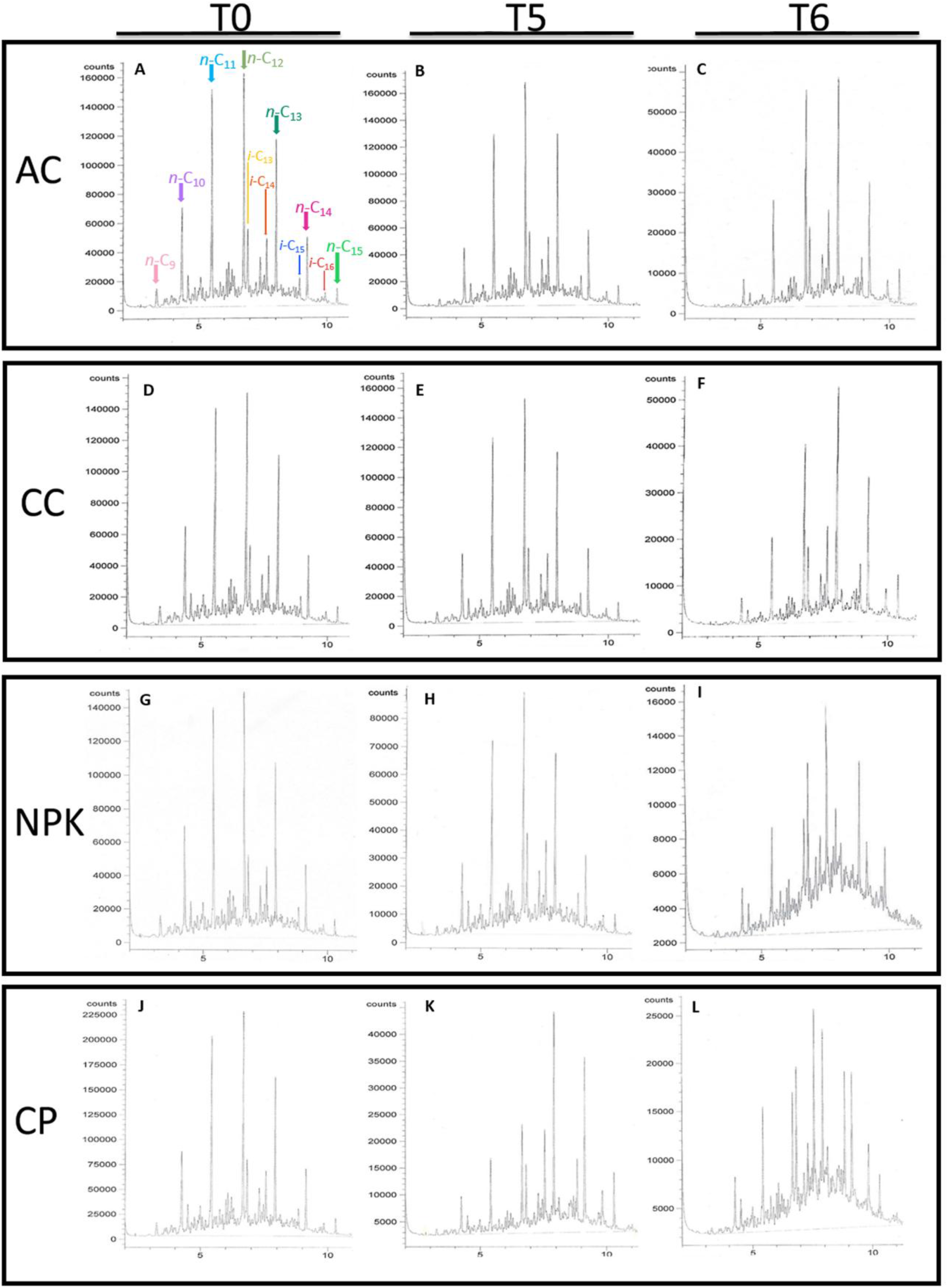
GC-FID chromatograms of the hydrocarbons extracted from soil samples. A-B-C: abiotic control (AC); D-E-F: non-fertilized control (CC); G-H-I: slow-release fertilizer Nitrofoska^®^ (NPK) and J-K-L: commercial product OSEII^®^ (CP) treatments. Samples were taken at day-0 (T0), day-50 (T5) and day-365 (T6) of the bioremediation experiment. Arrows indicate individual compounds.

### 3.3. Changes in the structure of bacterial communities

Bacterial communities were assessed using a culture-independent fingerprinting method. Comparing treatments, banding patterns of communities biostimulated with different nutrient sources were clearly different, as seen in the dendrograms constructed after the presence/absence analysis of DGGE bands (Fig. 5).

**Fig. 5:**
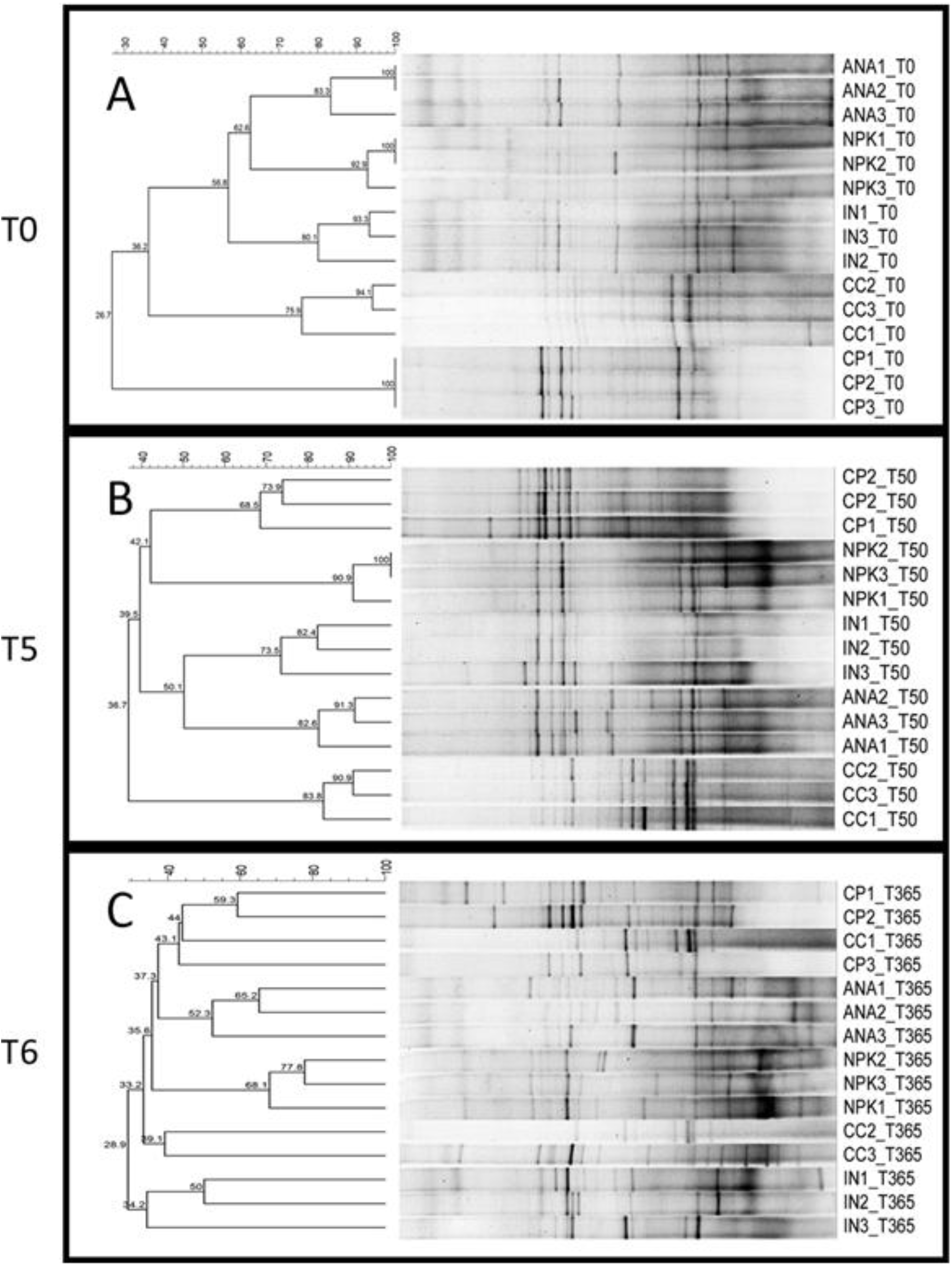
UPGMA dendrogram resulting from cluster analysis of data reporting the presence/absence of DGGE bands at day-0 (A), day-50 (B) and day-365 (C) of a bioremediation experiment with Antarctic soil chronically exposed to diesel fuel, using different biostimulation strategies. CC: non-fertilized control, IN: biostimulation with inorganics salts, ANA: biostimulation with inorganic salts without mixing, NPK: biostimulation with slow-release granular fertilizer, CP: biostimulation with a commercial bioremediation agent.

Although the soil used in the microcosms was thoroughly homogenized, the addition of different nutrients produced fast and differential changes in the structure of the native bacterial communities in that soil, as depicted by the differences in the DGGE profiles between CC and the other treatments even shortly after the preparation of the microcosms.

Bacterial community structure in CC changed after 50 days towards an enrichment in populations characterized by 16S rDNA sequences with mid/high GC content. These changes could be related to a positive effect of the soil aeration and mixing on the growth of some aerobic members of the indigenous microbiota, able to utilize or at least tolerate the hydrocarbons. Though different from CC, bacterial communities in IN and ANA microcosms after 50 days of treatment shared similar DGGE banding patterns, suggesting that aeration by mixing under biostimulation with inorganic salts did not produce substantial changes on the number and relative abundances of the populations detected by DGGE, at least under the assayed conditions. Nevertheless, after 365 days, IN and ANA communities diverged (28% similarity), even though no microcosm was mixed after day-50 and until the last sampling, in the following summer. The structures of bacterial communities present in NPK microcosms at the beginning of the experiment resembled those observed in IN and ANA, clustering together at 65% similarity. Towards day-30, the DGGE profiles were dominated by bands of low-GC% DNA sequences, already present since the beginning (Fig. S4), a trend that continued at least until day-50 (Fig. 5B). The most notorious and fast changes in bacterial community structure were observed in CP microcosms, clustering separately even at T0, where OSEII^®^ induced the enrichment of the indigenous microbiota in populations with low-GC% 16S rDNA represented by bands present in CP but not detected in the CC control. Over time, the intensity of the few bands already shared with CC at the beginning decreased, with a simultaneous increase in the intensity of other bands almost not detected in any other treatment, probably corresponding to bacteria favored only when nutrients and hydrocarbons are more readily assimilable due to the surfactants and enzymes supplied by OSEII^®^. Finally, as revealed by their similar DGGE banding patterns, the structures of the bacterial communities present in the independent replicate microcosms of each experimental treatment on day-0 were very similar to each other (Fig. S4), as was also observed at day-50. However, at day-365, DGGE profiles from replicates were not quite similar between them and exhibited multiple low-intensity bands spanning the entire gradient, likely depicting a higher richness and diversity than at day-50.

## 4. Discussion

Previous studies from our lab proved that autochthonous bacteria isolated from polluted soils at Carlini station have become adapted to the presence of hydrocarbons, acquiring the ability to metabolize these compounds (Ruberto et al., 2005; Vázquez et al., 2013). We also found that the bacterial growth and degradation efficiency were closely related with the concentration of N and P in the treated soil. (Dias et al., 2012, Martínez Álvarez et al., 2017). Our results here confirmed those observations, as biostimulation favored hydrocarbon biodegradation in chronically polluted soils from around Carlini Station. This enhanced degradation activity was accompanied by a significant increase in HDB counts throughout the experiment. In contrast, the low removal efficiency in the non-fertilized CC control was accompanied by low HAB and HDB counts, also proving that aeration alone was not enough to promote the enrichment of the soil microbiota in hydrocarbon degrading bacteria. Among the different nutrient sources tested for biostimulation of the indigenous microbiota, the commercial bioremediation agent OSEII^®^ and the slow-release fertilizer Nitrofoska^®^ promoted the fastest increases in bacterial numbers and hydrocarbon degradation rates, at short term and long term, respectively. Interestingly, although the increase in HAB numbers during the first ten days was similar in all the treatments including CC, HDB counts sharply increased in CP, NPK and IN. However, in NPK and IN the counts then dropped between days 10 and 20, whereas CP maintained a fairly constant and high value until the end of the experiment, which demonstrates that this bioremediation agent not only produces a rapid stimulation of the microbiota but also keeps it active throughout the process.

Concerning the fertilization with soluble inorganic salts, as they can be toxic at high concentrations due to osmotic effect (Walworth et al., 1997), we added them fractionally. In this sense, the periodic fluctuation in HDB counts, which was observed especially in the IN and ANA microcosm, could be reflecting a temporary lack of nutrients at adequate concentrations, caused by said fractional fertilization, to sustain a stable hydrocarbon degrading bacterial community. It is possible that, although the required amount of N and P to balance the C:N:P ratio in the soil to 100:10:1 was added, the strategy of dividing it in fractions representing 20% of the total amount has led to fluctuating C:N:P ratios, frequently differing from the optimum. Similarly, Dias et al. (2012) observed no significant differences in the efficiency of the biostimulated in comparison with the non-biostimulated microbiota in a 45-days long study performed in land plots, where the inorganic salts were added at once at the beginning of the experiment. However, we found biostimulation with inorganic salts successful in other experiments (Vázquez et al., 2009; Ruberto et al., 2010, Martínez Alvarez et al., 2017), which was also reported by other authors working with soils in cold environments (Margesin and Schinner, 2001; Powell et al., 2006). Clearly, when a bioremediation treatment is carried out outdoors and in open containers, the response of the soil microbiota is affected not only by the soil properties but also by the weather conditions, especially temperature, rain and snowfall. Fluctuations in bacterial counts could also have been the result of the soil freezing and thawing cycles or the precipitations, which cause the washing of the water dissolved salts. In any case, the fractionated fertilization did not seem to be successful in promoting a more efficient degradation of the hydrocarbons and, after 50 days of treatment, both IN and ANA microcosms showed no significant differences in degradation efficiency compared with the CC or even between them. Likewise, one year later, although IN and ANA reached a higher removal of hydrocarbons compared with CC, such differences were not statistically significant. This raises the question of whether it is possible in practice to maintain nutrient concentrations close to optimum by fertilizing with inorganic salts and, at the same time, avoid osmotic stress and washing, especially when working in open systems with polar mineral soils that often have low water holding capacity and experience periodic freezing and thawing, ice cover and snow melt during the summer (Delille and Coulon, 2008). In this sense, the biostimulation with slow-release fertilizers is one of the approaches used to overcome this problem, but the major challenge for their application resides in how to control the releasing rates to maintain optimal nutrient concentrations over long periods of time (Becker et al., 2016). We decided to evaluate Nitrofoska^®^ because it is cost effective and an efficient source of nutrients (Zhu et al., 2004), and it is promising to be used during the Antarctic winter, a long period where human intervention is not possible. In this study, Nitrofoska^®^ promoted the highest elimination of hydrocarbons (87%) after one year of treatment, evidencing that this product maintained its long-term fertilization capacity, sustaining the activity of the hydrocarbon degrading bacteria. The limited medium-term efficiency (day-50) of Nitrofoska^®^ could be due, to some extent, to the low soil temperatures that could have reduced the permeability of the slow-release particle coating (Lee et al., 1993), decreasing the nutrient release rate that could not thus meet the high demand of the large number of actively growing bacteria that develop in the early stages of the process.

Microbial communities from contaminated ecosystems had been described as less diverse than those inhabiting unstressed environments (Cury et al., 2015). Diversity is influenced by the complexity of chemical contaminants and by the time such populations have been exposed to the pollutants. In this regard, it should be kept in mind that the soil used here not only had a long history of exposure to hydrocarbons, but also suffered fresh contamination caused by frequent diesel leaks before being collected for this experiment, which makes it a complex and heterogeneous environment with many microhabitats suitable for different bacterial populations. In this scenario, not only well adapted oligotrophic microbes are established, but also opportunistic fast-growing copiotrophs can quickly react and play a relevant role in biodegradation (Martínez-Alonso et al., 2010). In this work, this was reflected by the differences in richness and diversity experienced by the bacterial communities along the process, also suggesting that the indigenous microbiota was able to rapidly react, playing an active role in the removal of hydrocarbons. In particular, OSEII^®^ (recommended by EPA for bioremediation in cold areas) promoted a rapid and stable response of the microbiota, with a sharp increase in HDB numbers shortly after addition, and favored bacterial populations that led to a significantly higher hydrocarbon removal than the achieved with the other strategies, after only 50 days of treatment. This quick response of the microbiota would be represented by the DGGE bands with increased relative abundance observed in the soil samples taken from CP microcosms at day-50. Using the same commercial product in land plots with chronically contaminated soil at Carlini Station, Dias et al. (2012) also found a significantly higher hydrocarbon removal, but reported minor differences in the DGGE patterns relative to the non-biostimulated microbiota.

It is well known that diesel fuel components differ in their susceptibility to microbial attack (Chikere et al., 2011), and that the lightest fractions are biodegradable but also volatile and more susceptible to abiotic loss. Other compounds, like the isoprenoids, are more recalcitrant to biological degradation due to their branched chemical structure, although their degradation under natural conditions was proved and the corresponding catabolic pathways are known (Varjani, 2017). Gas chromatography profiles in this experiment revealed a significant loss by volatilization and washing, which was the main cause of hydrocarbon removal in the non-biostimulated microcosms. As all microcosms had the same geometry and experimental design, similar abiotic loss was expected for CC and the fertilized microcosms and, therefore, the higher hydrocarbon removal reached by the fertilized microcosms could be attributed to the activity of the microbiota. In addition, the chromatograms obtained from soil samples at the beginning and after 50 and 365 days of treatment showed that the degradation of the hydrocarbons was lower the longer their carbon chain, and that the linear alkanes degraded more readily than their corresponding isoprenoids of similar carbon chain length. The same was reported to occur also during the bioremediation of other contaminated soils around the world (Aislabie et al., 2006; Coulon and Delille, 2006). In this sense, relative amounts of certain compounds could be indicative of the extent of biodegradation versus abiotic loss for complex mixtures of hydrocarbons. According to the model proposed by Snape et al. 2005, when biodegradation can be excluded as a process, the ratio of *i*-C13/*i*-C16 is a sensitive measure of evaporation over a wide range of intermediate to major mass loss, while the ratios of *n*-C9-12/*i*-C16 would be better indicators of minor loss of mass by evaporation. In the abiotic control, biodegradation can be excluded and, therefore, it can be assumed that the loss of hydrocarbons was produced by washing (without major changes in the proportion of the compounds in the remaining AGO) and by evaporation (producing the fractionation of the most volatile compounds). In a biodegradation experiment without loss by evaporation, Snape et al. 2005 found that ratios such as *n*-C12/*i*-C13 and *n*-C13/*i*-C14 decreased in comparison with abiotic controls and the isoprenoids did not change throughout the experiment, while the n-alkane/isoprenoid ratios of similar volatilities changed as biodegradation proceeded. By comparing these ratios in the CC and AC controls at each time interval (T0-T5, T5-T6) it would seem that the light hydrocarbons in CC were mainly lost by lixiviation and evaporation, with some biodegradation at the end of the experiment. Conversely, in NPK after one year and in CP already at day-50, *n*-C12/*i*-C13 and *n*-C13/*i*-C14 decreased considerably to half or less the values in AC, indicating that the changes observed were also of biological origin. This supports our observation about the effectiveness of CP to induce the necessary changes in the microbial community to promote biodegradation in a short time, resulting in an efficient bioremediation treatment.

As mentioned in the results section, after one year of treatment the initial concentration of hydrocarbons in all the non-poisoned microcosms was reduced by about 50%. This suggests that the HDB likely remained active and continued with their catabolic activity during winter, despite the very low temperature of the frozen soil, probably inhabiting microenvironments where the water remained liquid. This was supported by the DGGE banding patterns obtained from all the biostimulated treatments at day-365, that showed notorious and distinct changes in their community structures, with higher richness and evenness compared to the original community. A similar increase in bacterial diversity at the end of a successful bioremediation experiment was also reported by Dias et al. (2012) working with soils from Carlini Station. In our work, we found similar DGGE banding patterns associated with independent replicates from the same treatment at the beginning of the experiment and at day-50. This suggests that, during the first 50 days of treatment, the dynamics followed by the soil bacterial microbiota was mainly driven by the applied biostimulation strategy more than by the environmental factors, although the replicate microcosms frequently differed in terms of water content or presence of frozen soil or snow cover. Banding patterns from DGGE are not always considered accurate enough to calculate richness and diversity indices (Neilson et al., 2013). However, the high reproducibility obtained with our soil samples allowed us to use the presence/absence of shared bands to build a dendrogram and report on the main differences between treatments regarding the evolution of bacterial communities, as was also done by others (Schauer et al., 2000; Gafan et al., 2005; Chong et al., 2009; Bevivino et al., 2014; Festa et al., 2016).

It is possible that the different dynamics followed by the bacterial communities under the different biostimulation strategies assayed were related to the gradual reduction of environmental stress that the hydrocarbon contamination represented for the indigenous microbiota, which occurred at different rates and extent according to the applied strategy. Once the biostimulation agents were added, the balance of nutrients under such stress could have triggered the development of the fast-growing hydrocarbon-tolerant members of the microbiota, capable to efficiently catabolize the easily degradable fraction of the contaminants at the early stages of the process. This could have precluded the detection by DGGE of other more sensitive or slow-growing hydrocarbon degraders that would be present in the soil but at very low proportions. In this sense, while DGGE fingerprinting at day-50 revealed rather similar bacterial community structures between the replicate microcosms from a single treatment, the patterns differed between treatments. All fertilized microcosms showed high abundance of bacteria with medium to low-GC% 16S rDNA, compatible with fast hydrocarbon-degraders like *Pseudomonas* and other Proteobacteria. Conversely, non-fertilized microcosms (CC) showed DGGE patterns dominated by high-GC% 16S rDNA bands, compatible with *Actinobacteria* and other more oligotrophic and slow-growing bacteria, adapted to resource-limited conditions. As mentioned above, after 365 days of treatment the bacterial communities diverged in a way that they were different even in the replicate microcosms. In this sense, Yan et al. (2016) postulated that a successional variation in the composition of the bacterial community represents a sensitive ecological indicator of *in situ* remediation, and White et al. (1998) stated that pollution disappearance accompanied by consequent changes in the structure of microbial communities is indicative of a reduced stress in the microniche environment influencing the local microbiota, even if there has not been a return to the baseline community. After one year, the biological degradation of the organic compounds leading to a depletion of the amount of nutrients in the biostimulated microcosms may have promoted the establishment of bacterial communities dominated by *k*-strategists, able to cope with a reduced availability of resources.

## 5. Conclusion

Although more tests are needed, it would appear that the commercial product used in the CP microcosms (a mixture of surfactants, nutrients and enzymes) determines a better mid-term performance while NPK (a slow-release fertilizer), which prevailed as the best source of nutrients one year after its addition, is more efficient for long-term processes. Our findings suggest that a mixed strategy combining the fastest action of commercial products like OSEII^®^ during the Antarctic summer with slow-release fertilizers like Nitrofoska^®^, more active during the rest of the year, could be an efficient long-term bioremediation treatment for Antarctic areas where the temperature rises above the freezing point and the ground is free of ice and snow for a short time during the summer. To conclude, understanding how the species interactions, nutrients, and changes in quantity and bioavailability of hydrocarbons affect the structure and degradation capability of a microbial community represents a key factor to improve the design of proper and customized bioremediation strategies, highlighting the importance of conducting research in this topic to remediate Antarctic soils.

## 6. Acknowledgments

This work was supported by the Agencia Nacional de Promoción Científica y Tecnológica, ANPCyT (PICT 2016-7271) and the Universidad de Buenos Aires, UBA (UBACyT 20020130100569BA). We are grateful to the Instituto Antártico Argentino (IAA), Carlini Station, for their hospitality and logistic support and to Silvia H. Coria for her valuable support in the field and laboratory activities performed at Carlini Station.

## References

Abed, R.M., Al-Sabahi, J., Al-Maqrashi, F., Al-Habsi, A., Al-Hinai, M., 2014. Characterization of hydrocarbon-degrading bacteria isolated from oil-contaminated sediments in the Sultanate of Oman and evaluation of bioaugmentation and biostimulation approaches in microcosm experiments. Int. Biodeterior. Biodegradation 89, 58–66. https://doi.org/10.1016/j.ibiod.2014.01.006

Aislabie, J.M., Balks, M.R., Foght, J.M., Waterhouse, E.J., 2004. Hydrocarbon Spills on Antarctic Soils: Effects and Management. Environ. Sci. Technol. 38, 1265–1274. https://doi.org/10.1021/es0305149

Aislabie, J., Saul, D.J., Foght, J.M., 2006. Bioremediation of hydrocarbon-contaminated polar soils. Extremophiles 10, 171–179. https://doi.org/10.1007/s00792-005-0498-4

Becker, R.B., de Souza, E.S., Martins, R.L., da Luz Bueno, J., 2016. Bioremediation of Oil-Contaminated Beach and Restinga Sediments Using a Slow-Release Fertilizer. Clean Soil Air Water. 44, 1154–1162. https://doi.org/10.1002/clen.201500023

Bennett, J.R., Shaw, J.D., Terauds, A., Smol, J.P., Aerts, R., Bergstrom, D.M., Blais, J.M., Cheung, W.W.L., Chown, S.L., Lea, M.A., Nielsen, U.N., Pauly, D., Reimer, K.J., Riddle, M.J., Snape, I., Stark, J.S., Tulloch, V.J., Possingham, H.P., 2015. Polar lessons learned: long-term management based on shared threats in Arctic and Antarctic environments. Front Ecol Env. 13, 316–324. https://doi.org/10.1890/140315

Bevivino, A., Paganin, P., Bacci, G., Florio, A., Pellicer, M.S., Papaleo, M.C., Mengoni, A., Ledda, L., Fani, R., Benedetti, A., Dalmastri, C., 2014. Soil Bacterial Community Response to Differences in Agricultural Management along with Seasonal Changes in a Mediterranean Region. PLoS ONE. 9, e105515. https://doi.org/10.1371/journal.pone.0105515

Braun, C., Hertel, F., Mustafa, O., Nordt, A., Pfeiffer, S., Peter, H.U., 2014. Environmental Assessment and Management Challenges of the Fildes Peninsula Region. In: Tin T., Liggett D., Maher P., Lamers M. (eds) Antarctic Futures. Springer, Dordrecht. Chapter 1. https://doi.org/10.1007/978-94-007-6582-5_7

Camenzuli, D., Freidman, B.L., 2015. On-site and in situ remediation technologies applicable to petroleum hydrocarbon contaminated sites in the Antarctic and Arctic Antarctic and Arctic. Polar Res. 34, 24492. https://doi.org/10.3402/polar.v34.24492

Chikere, C.B., Okpokwasili, G.C., Chikere, B.O., 2011. Monitoring of microbial hydrocarbon remediation in the soil. 3 Biotech 1, 117–138. https://doi.org/10.1007/s13205-011-0014-8

Chong, C.W., Annie Tan, G.Y., Wong, R.C.S., Riddle, M.J., Tan, I.K.P., 2009. DGGE fingerprinting of bacteria in soils from eight ecologically different sites around Casey Station, Antarctica. Polar Biol. 32, 853–860. https://doi.org/10.1007/s00300-009-0585-6

Coulon, F., Delille, D., 2006. Influence of substratum on the degradation processes in diesel polluted sub-Antarctic soils (Crozet Archipelago). Polar Biol. 29, 806–812. https://doi.org/10.1007/s00300-006-0118-5

Cury, J.C., Jurelevicius, D.A., Villela, H.D.M., Jesus, H.E., Peixoto, R.S., Schaefer, C.E.G.R., Bícego, M.C., Seldin, L., Rosado, A.S., 2015. Microbial diversity and hydrocarbon depletion in low and high diesel-polluted soil samples from Keller Peninsula, South Shetland Islands. Antarct. Sci. 27, 263–273. https://doi.org/10.1017/S0954102014000728

Dadrasnia, A., Agamuthu, P., 2014. Biostimulation and monitoring of diesel fuel polluted soil amended with biowaste. Pet. Sci. Technol. 32, 2822–2828. https://doi.org/10.1080/10916466.2014.913624

de Jesus, H.E., Peixoto, R.S., Rosado, A.S., 2015. Bioremediation in Antarctic Soils. J Pet Env. Biotechnol 6:248. https://doi.org/10.4172/2157-7463.1000248

Delille, D., Coulon, F., 2008. Comparative Mesocosm Study of Biostimulation Efficiency in Two Different Oil-Amended Sub-Antarctic Soils. Microb Ecol. 56, 243–252. https://doi.org/10.1007/s00248-007-9341-z

Dias, R.L., Ruberto, L., Hernández, E.A., Vázquez, S.C., Lo Balbo, A., Del Panno, M.T., Mac Cormack, W.P., 2012. Bioremediation of an aged diesel oil-contaminated Antarctic soil: Evaluation of the “on site” biostimulation strategy using different nutrient sources. Int. Biodeterior. Biodegrad. 75, 96–103. https://doi.org/10.1016/j.ibiod.2012.07.020

Dias, R.L., Ruberto, L., Calabró, A., Lo Balbo, A., Del Panno, M.T., Mac Cormack, W.P., 2015. Hydrocarbon removal and bacterial community structure in on-site biostimulated biopile systems designed for bioremediation of diesel-contaminated Antarctic soil. Polar Biol. 38, 677–687. https://doi.org/10.1007/s00300-014-1630-7

Emami, S., Pourbabaee, A.A., Alikhani, H.A., 2014. Interactive effect of nitrogen fertilizer and hydrocarbon pollution on soil biological indicators. Environ. Earth Sci. 72, 3513–3519. https://doi.org/10.1007/s12665-014-3259-9

EPA, 1999. Monitored natural attenuation of petroleum hydrocarbons. Report EPA/600/F-98/021, Office of Research and Development, Environmental Protection Agency. U.S, Washington, D.C.

Ezenne, G.I., Nwoke, O.A., Ezikpe, D.E., Obalum, S.E., Ugwuishiwu, B.O., 2014. Use of poultry droppings for remediation of crude-oil-polluted soils: Effects of application rate on total and poly-aromatic hydrocarbon concentrations. Int. Biodeterior. Biodegrad. 92, 57–65. https://doi.org/10.1016/j.ibiod.2014.01.025

Festa, S., Coppotelli, B.M., Morelli, I.S., 2016. Comparative bioaugmentation with a consortium and a single strain in a phenanthrene-contaminated soil: Impact on the bacterial community and biodegradation. Appl. Soil Ecol. 98, 8–19. https://doi.org/10.1016/J.APSOIL.2015.08.025

Gafan, G.P., Lucas, V.S., Roberts, G.J., Petrie, A., Wilson, M., Spratt, D.A., 2005. Statistical Analyses of Complex Denaturing Gradient Gel Electrophoresis Profiles. J. Clin. Microbiol. 43, 3971–3978. https://doi.org/10.1128/JCM.43.8.3971-3978.2005

Goswami, M., Chakraborty, P., Mukherjee, K., Mitra, G., Bhattacharyya, P., Dey, S., Tribedi, P., 2018. Bioaugmentation and biostimulation: a potential strategy for environmental remediation. J. Microbiol. Exp. 6, 223–231. https://doi.org/10.15406/jmen.2018.06.00219

Grace Liu, P.W., Chang, T.C., Whang, L.M., Kao, C.H., Pan, P.T., Cheng, S.S., 2011. Bioremediation of petroleum hydrocarbon contaminated soil: Effects of strategies and microbial community shift. Int. Biodeterior. Biodegrad. 65, 1119–1127. https://doi.org/10.1016/j.ibiod.2011.09.002

Hamzah, A., Phan, C.W., Yong, P.H., Mohd Ridzuan, H.N., 2014. Oil Palm Empty Fruit Bunch and Sugarcane Bagasse Enhance the Bioremediation of Soil Artificially Polluted by Crude Oil. Soil Sediment Contam. An Int. J. 23, 751–762. https://doi.org/10.1080/15320383.2014.870528

Huesemann, M., 2004. Biodegradation and Bioremediation of Petroleum Pollutants in Soil. In: Singh A., Ward O.P. (eds) Applied Bioremediation and Phytoremediation. Soil Biology. Vol. 1. Springer, Berlin, Heidelberg. https://doi.org/10.1007/978-3-662-05794-0_2

Hughes, K.A., Stallwood, B., 2006. Oil Pollution in the Antarctic Terrestrial Environment. Polarforshung 75, 141–144.

Kachinskii, V.L., Zavgorodnyaya, Y.A., Gennadiev, A.N., 2014. Hydrocarbon contamination of Arctic Tundra Soils of the Bol’shoi Lyakhovskii Island (the Novosibirskie Islands). Eurasian Soil Sci. 47, 57–69. https://doi.org/10.1134/S1064229314020070

Kauppi, S., Sinkkonen, A., Romantschuk, M., 2011. Enhancing bioremediation of diesel-fuel-contaminated soil in a boreal climate: Comparison of biostimulation and bioaugmentation. Int. Biodeterior. Biodegrad. 65, 359–368. https://doi.org/10.1016/j.ibiod.2010.10.011

Lee, K., Tremblay, G.H., Levy, E.M., 1993. Bioremediation: Application of Slow-Release Fertilizers on Low-Energy Shorelines. Int. Oil Spill Conf. Proc. 1993, 449–454. https://doi.org/10.7901/2169-3358-1993-1-449

Lee, S.H., Lee, S., Kim, D., Kim, J., 2007. Degradation characteristics of waste lubricants under different nutrient conditions. J. Hazard. Mater. 143, 65–72. https://doi.org/10.1016/j.jhazmat.2006.08.059

Margesin, R., Schinner, F., 2001. Biodegradation and bioremediation of hydrocarbons in extreme environments. Appl. Microbiol. Biotechnol. 56, 650–663. https://doi.org/10.1007/s002530100701

Martínez Álvarez, L., Ruberto, L., Lo Balbo, A., Mac Cormack, W.P, 2017. Bioremediation of hydrocarbon-contaminated soils in cold regions: Development of a pre-optimized biostimulation biopile-scale field assay in Antarctica. Sci. Total Environ. 590-591, 194–203. https://doi.org/10.1016/j.scitotenv.2017.02.204

Martínez-Alonso, M., Escolano, J., Montesinos, E., Gaju, N., 2010. Diversity of the bacterial community in the surface soil of a pear orchard based on 16s rRNA gene analysis. Int. Microbiol. 13, 123–134. https://doi.org/10.2436/20.1501.01.117

Mulligan, C.N., Yong, R.N., 2004. Natural attenuation of contaminated soils. Environ. Int. 30, 587–601. https://doi.org/10.1016/j.envint.2003.11.001

Muyzer, G., De Waal, E.C., Uitterlinden, A.G., 1993. Profiling of Complex Microbial Populations by Denaturing Gradient Gel Electrophoresis Analysis of Polymerase Chain Reaction-Amplified Genes Coding for 16S rRNA. Appl. Environ. Microbiology 59, 695–700. PubMed PMID: 7683183; PubMed Central PMCID: PMC202176.

Neilson, J.W., Jordan, F.L., Maier, R.M., 2013. Analysis of Artifacts Suggests DGGE Should Not Be Used For Quantitative Diversity Analysis. J. Microbiol Methods 92, 256–263. https://doi.org/10.1016/j.mimet.2012.12.021

Powell, S.M., Ferguson, S.H., Snape, I., Siciliano, S.D., 2006. Fertilization stimulates anaerobic fuel degradation of antarctic soils by denitrifying microorganisms. Environ. Sci. Technol. 40, 2011–2017. https://doi.org/10.1021/es051818t

Raymond, T., King, C.K., Raymond, B., Stark, J.S., Snape, I., 2017. Oil Pollution in Antarctica. In: Mervin Fingas (eds) Oil Spill Science and Technology (Second Edition). Gulf Professional Publishing. Chapter 14. 759–803. https://doi.org/10.1016/B978-0-12-809413-6.00014-X

Rittmann, B.E., Hausner, M., Löffler, F., Love, N.G., Muyzer, G., Okabe, S., Oerther, D.B., Peccia, J., Raskin, L., Wagner, M., 2006. A Vista for Microbial Ecology and Environmental Biotechnology. Environ. Sci. Technol. 40, 1096–1103. https://doi.org/10.1021/es062631k

Ruberto, L., Vazquez, S.C., Mac Cormack, W.P., 2003. Effectiveness of the natural bacterial flora, biostimulation and bioaugmentation on the bioremediation of a hydrocarbon contaminated Antarctic soil. Int. Biodeterior. Biodegrad. 52, 115–125. https://doi.org/10.1016/S0964-8305(03)00048-9

Ruberto, L., Vazquez, S.C., Lo Balbo, A., Mac Cormack, W.P, 2005. Psychrotolerant hydrocarbon-degrading Rhodococcus strains isolated from polluted Antarctic soils. Antarct. Sci. 17, 47–56. https://doi.org/10.1017/S0954102005002415

Ruberto, L., Dias, R.L., Lo Balbo, A., Vazquez, S.C., Hernandez, E.A., Mac Cormack, W.P., 2009. Influence of nutrients addition and bioaugmentation on the hydrocarbon biodegradation of a chronically contaminated Antarctic soil. J. Appl. Microbiol. 106, 1101–1110. https://doi.org/10.1111/j.1365-2672.2008.04073.x

Ruberto, L., Vazquez, S.C, Dias, R.L., Hernández, E.A., Coria, S.H., Levin, G., Lo Balbo, A., Mac Cormack, W.P. 2010. Small-scale studies towards a rational use of bioaugmentation in an Antarctic hydrocarbon-contaminated soil. Antarct. Sci. 22, 463/469. https://doi.org/10.1017/S0954102010000295

Sampaio, D.S., Almeida, J.R.B., de Jesus, H.E., Rosado, A.S., Seldin, L., Jurelevicius, D., 2017. Distribution of Anaerobic Hydrocarbon-Degrading Bacteria in Soils from King George Island, Maritime Antarctica. Microb. Ecol. 74, 810–820. https://doi.org/10.1007/s00248-017-0973-3

Schauer, M., Massana, R., Pedrós-Alió, C., 2000. Spatial differences in bacterioplankton composition along the Catalan coast (NW Mediterranean) assessed by molecular fingerprinting. FEMS Microbiol. Ecol. 33, 51–59. https://doi.org/10.1111/j.1574-6941.2000.tb00726.x

Silva-Castro, G.A., Uad, I., Rodríguez-Calvo, A., González-López, J., Calvo, C., 2015. Response of autochthonous microbiota of diesel polluted soils to land-farming treatments. Environ. Res. 137, 49–58. https://doi.org/10.1016/j.envres.2014.11.009

Simpanen, S., Dahl, M., Gerlach, M., Mikkonen, A., Malk, V., Mikola, J., Romantschuk, M., 2016. Biostimulation proved to be the most efficient method in the comparison of in situ soil remediation treatments after a simulated oil spill accident. Environ. Sci. Pollut. Res. 23, 25024–25038. https://doi.org/10.1007/s11356-016-7606-0

Snape, I., McA Harvey, P., Ferguson, S.H., Rayner, J.L., Revill, A.T., 2005. Investigation of evaporation and biodegradation of fuel spills in Antarctica I. A chemical approach using GC – FID. Chemosphere. 61, 1485–1494. doi:10.1016/j.chemosphere.2005.04.108

Szopińska, M., Namieśnik, J., Polkowska, Ż., 2016. How Important Is Research on Pollution Levels in Antarctica? Historical Approach, Difficulties and Current Trends. In: de Voogt P. (eds) Reviews of Environmental Contamination and Toxicology. Reviews of Environmental Contamination and Toxicology (Continuation of Residue Reviews). Springer, Cham. 239, 79–156. https://doi.org/10.1007/398_2015_5008

Varjani, S., 2017. Microbial degradation of petroleum hydrocarbons. Bioresource Technology, 223, 277–286. https://doi.org/10.1016/j.biortech.2016.10.037

Vázquez, S.C, Nogales, B., Ruberto, L., Hernández, E.A, Christie-Oleza, J., Lo Balbo, A., Bosch, R., Lalucat, J., Mac Cormack, W.P, 2009. Bacterial community dynamics during bioremediation of diesel oil-contaminated antarctic soil. Microb. Ecol. 57, 598–610. https://doi.org/10.1007/s00248-008-9420-9

Vázquez, S.C, Nogales, B., Ruberto, L., Mestre, C., Christie-Oleza, J., Ferrero, M., Bosch, R., Mac Cormack, W.P., 2013. Characterization of bacterial consortia from diesel-contaminated Antarctic soils: Towards the design of tailored formulas for bioaugmentation. Int. Biodeterior. Biodegrad. 77, 22–30. https://doi.org/10.1016/j.ibiod.2012.11.002

Vázquez, S.C., Monien, P., Pepino Minetti, R., Jürgens, J., Curtosi, A., Villalba Primitz, J., Frickenhaus, S., Abele, D., Mac Cormack, W.P., Helmke, E., 2017. Bacterial communities and chemical parameters in soils and coastal sediments in response to diesel spills at Carlini Station, Antarctica. Sci. Total Environ. 605-606, 26–37. https://doi.org/10.1016/J.SCITOTENV.2017.06.129

Walworth, J.L., Woolard, C.R., Braddock, J.F., Reynolds, C.M., 1997. Enhancement and inhibition of soil petroleum biodegradation through the use of fertilizer nitrogen: An approach to determining optimum levels. J. Soil Sediment Contam. 6, 465–480. https://doi.org/10.1080/15320389709383580

White, D.C., Flemming, C.A., Leung, K.T., MacNaughton, S.J., 1998. In situ microbial ecology for quantitative appraisal, monitoring, and risk assessment of pollution remediation in soils, the subsurface, the rhizosphere and in biofilms. J. Microbiol. Methods 32, 93–105. https://doi.org/10.1016/S0167-7012(98)00017-7

Wu, M., Dick, W.A., Li, W., Wang, X., Yang, Q., Wang, T., Xu, L., Zhang, M., Chen, L., 2016. Bioaugmentation and biostimulation of hydrocarbon degradation and the microbial community in a petroleum-contaminated soil. Int. Biodeterior. Biodegrad. 107, 158–164. https://doi.org/10.1016/j.ibiod.2015.11.019

Yan, L., Sinkko, H., Penttinen, P., Lindström, K., 2016. Characterization of successional changes in bacterial community composition during bioremediation of used motor oil-contaminated soil in a boreal climate. Sci. Total Environ. 542, 817–825. https://doi.org/10.1016/j.scitotenv.2015.10.144

Zawierucha, I., Malina, G., 2011. Bioremediation of Contaminated Soils: Effects of Bioaugmentation and Biostimulation on Enhancing Biodegradation of Oil Hydrocarbons. In: Singh A., Parmar N., Kuhad R. (eds) Bioaugmentation, Biostimulation and Biocontrol. Soil Biology. Springer, Berlin, Heidelberg. vol 108. Chapter 1. https://doi.org/10.1007/978-3-642-19769-7_8

Zhong, W., An, H., Fang, W., Gao, X., Dong, D., 2016. Features and evolution of international fossil fuel trade network based on value of emergy. Appl. Energy 165, 868–877. https://doi.org/10.1016/j.apenergy.2015.12.083

Zhu, X., Venosa, A.D., Suidan, M.T., 2004. Literature Review on the Use of Commercial Bioremediation Agents for Cleanup of Oil-Contaminated Estuarine Environments. Report Epa/600/R-04/075. National Risk Management Research Laboratory, Office of Research and Development, Environmental Protection Agency. U.S, Cincinnati. pp. 1–56.

