## Supplementary information for "Bioremediation of hydrocarbon contaminated soil from Carlini Station, Antarctica: effectiveness of different nutrient sources as biostimulation agents"

Supplementary Figure S1: Experimental design

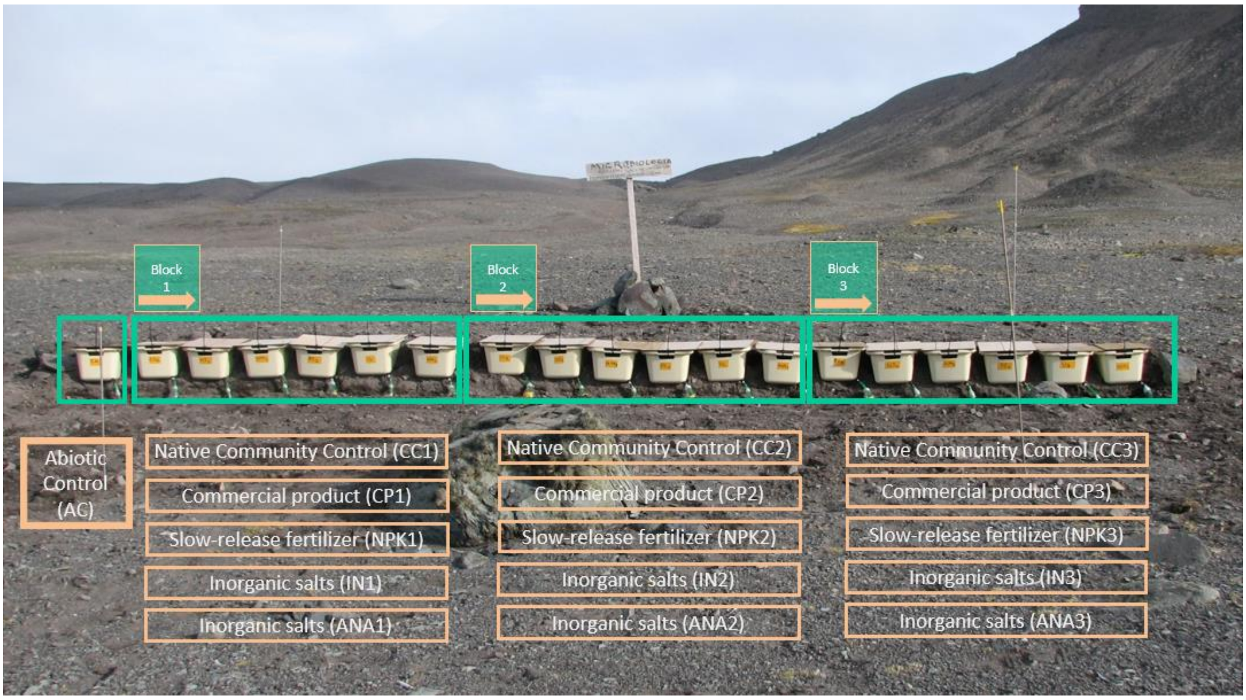


The microcosms were distributed in the soil in three blocks, each block containing one of the three replicates of the biostimulation treatments (IN, ANA, NPK, CP) and the non-fertilized control (CC), to make the influence of the different impact of winds, rains, snowfall and temperature fluctuations be more homogeneous between treatments.

Supplementary Figure S2: Sampling strategy

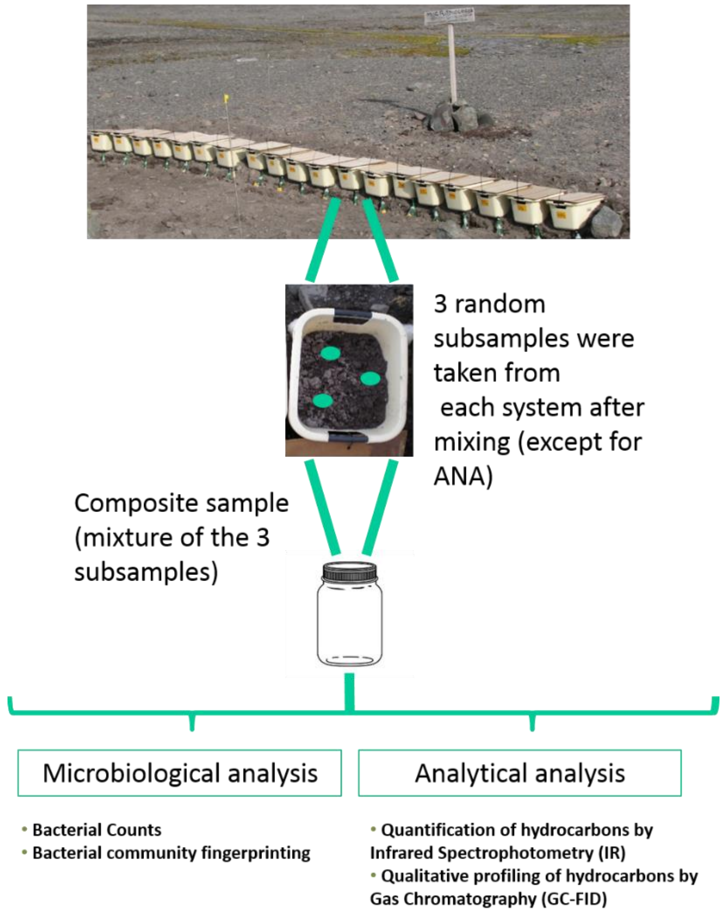


Supplementary Figure S3: GC-FID chromatogram profile of the diesel fuel (Antarctic gasoil: AGO) used at Carlini Station, showing the main lineal alkanes and isoprenoids.

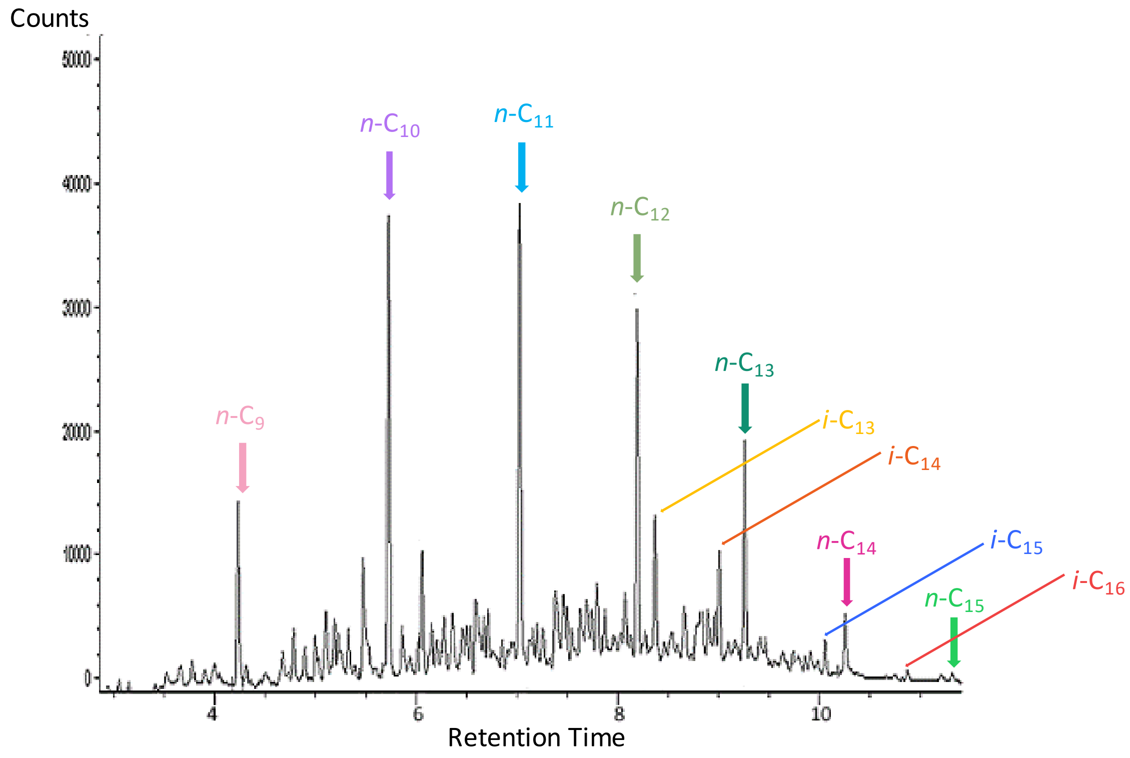

Supplementary Figure S4: DGGE fingerprinting of bacterial communities present in the all soil samples taken throughout the bioremediation experiment.

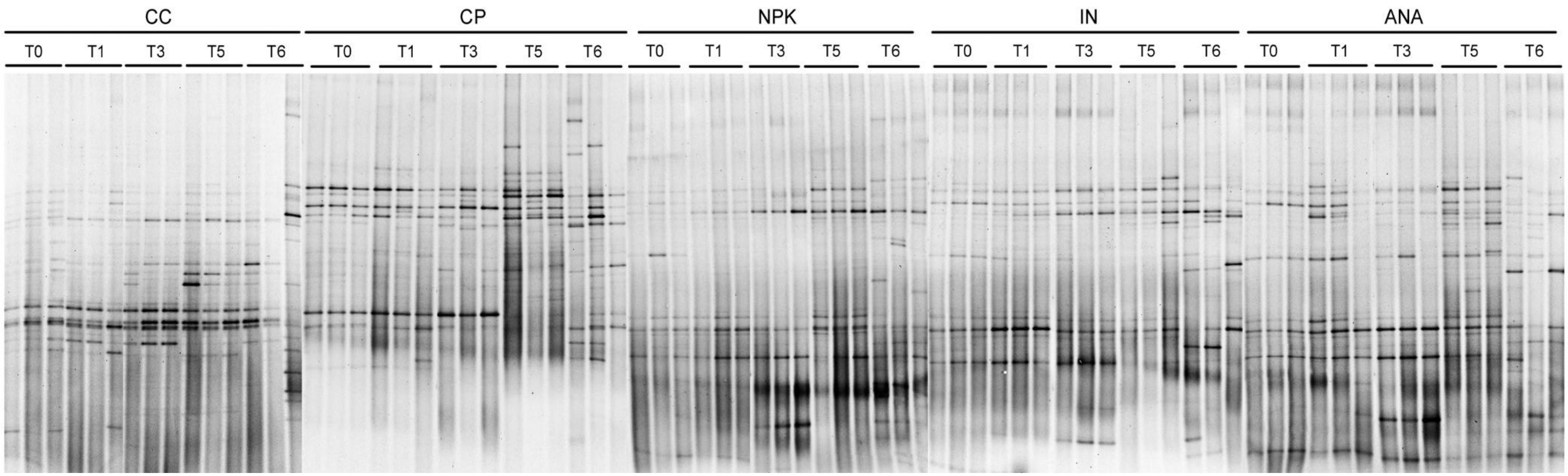
